# Atomistic Insights into gp82 Binding: A Microsecond, Million-Atom Exploration of Trypanosoma cruzi Host-Cell Invasion

**DOI:** 10.1101/2024.10.22.619626

**Authors:** Raissa S. L. Rosa, Manuela Leal da Silva, Rafael C. Bernardi

**Affiliations:** Department of Physics, Auburn University, Auburn, AL, USA; Department of Chemistry and Biochemistry, Auburn University, Auburn, AL, USA; Programa de Pos Graduacao em Biologia Computacional e Sistema, Instituto Oswaldo Cruz (FIOCRUZ), Rio de Janeiro, Brazil; Instituto de Biodiversidade e Sustentabilidade (NUPEM), Universidade Federal do Rio de Janeiro, Macaé, Brazil

## Abstract

Chagas disease, caused by the protozoan *Trypanosoma cruzi*, affects millions globally, leading to severe cardiac and gastrointestinal complications in its chronic phase. The invasion of host cells by *T. cruzi* is mediated by the interaction between the parasite’s glycoprotein gp82 and the human receptor lysosome-associated membrane protein 2 (LAMP2). While experimental studies have identified a few residues involved in this interaction, a comprehensive molecular-level understanding has been lacking. In this study, we present a 1.44-million-atom computational model of the gp82 complex, including over 3,300 lipids, glycosylation sites, and full molecular representations of gp82 and LAMP2, making it the most complete model of a parasite-host interaction to date. Using microsecond-long molecular dynamics simulations and dynamic network analysis, we identified critical residue interactions, including novel regions of contact that were previously uncharacterized. Our findings also highlight the significance of the transmembrane domain of LAMP2 in stabilizing the complex. These insights extend beyond traditional hydrogen bond interactions, revealing a complex network of cooperative motions that facilitate *T. cruzi* invasion. This study not only confirms key experimental observations but also uncovers new molecular targets for therapeutic intervention, offering a potential pathway to disrupt *T. cruzi* infection and combat Chagas disease.

## Introduction

Chagas disease (CD), caused by the parasitic protozoan *Trypanosoma cruzi*, is endemic in Latin America and primarily transmitted by triatomine bugs, which are insects mainly found in the Americas.^1,2^ These blood-sucking insects typically live in the walls and roofs of homes in rural and suburban areas and emerge at night to feed on human and animal hosts. Historically isolated in Latin America, the global impact of CD has escalated due to increased international migration, which has introduced the parasite to non-endemic regions.^3–5^ Transmission can also occur through congenital means during pregnancy, blood transfusions, organ transplants, and, less commonly, through contaminated food or drinks. ^6^ According to the World Health Organization (WHO), the disease affects an estimated 6 to 7 million people worldwide, with approximately 30,000 new cases reported annually, including 8,000 cases in newborns who acquire the infection during pregnancy. Furthermore, about 70 million individuals are at risk of contracting the disease due to their residence in areas of potential exposure.^6^ CD, though curable if treated early in the acute phase, can lead to severe cardiac, digestive, and neurological complications in its chronic phase, especially if diagnosis and treatment are delayed.^7^ The disease is often di”cult to diagnose in its early stages due to the presence of multiple non-specific symptoms, causing many cases to progress to the chronic phase. ^8^

The infection life cycle of *T. cruzi* begins when the triatomine vector releases metacyclic trypomastigotes (MTs) onto the host’s skin during a bite (see **Fig.1a**).^4^ Once in the blood-stream, MTs migrate to various tissues, invade nucleated cells, and differentiate and multiply in the cytoplasm.^5^ To facilitate entry and survival, *T. cruzi* exploits host signaling pathways, with calcium (Ca^2+^) playing a crucial role in these processes. ^9^ Typically, the host’s lysosomal machinery aids in membrane repair, recycling, and immune defense by fusing with damaged membranes and degrading foreign invaders.^10^ However, *T. cruzi* evades this by preventing phagolysosome maturation and acidification,^11^ and before lysosomal destruction can occur, the parasite escapes into the cytoplasm where it safely replicates. ^12^

**Figure 1:**
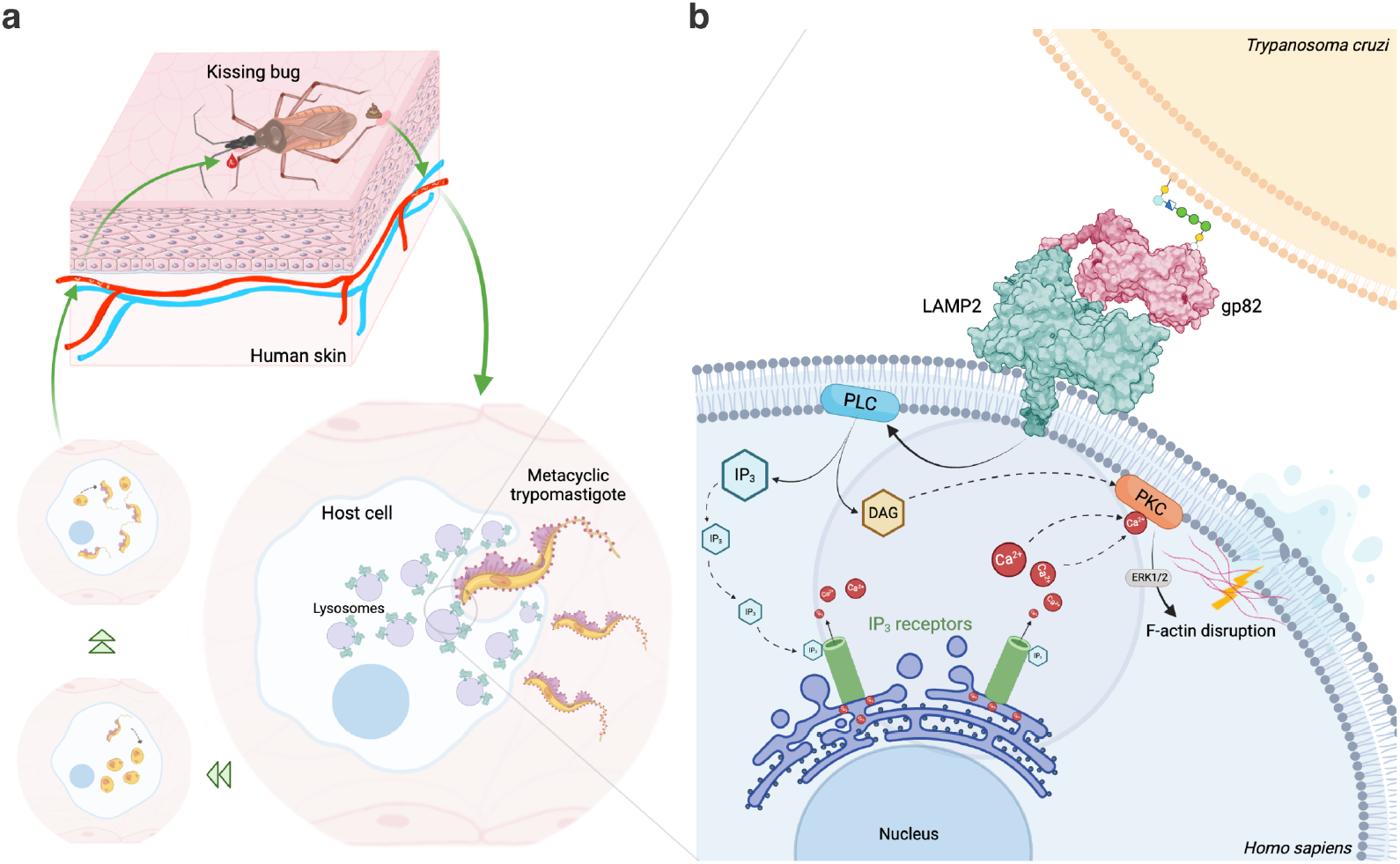
Host life cycle and infection interface. (a) *T. cruzi* life cycle: During the feeding process, the insect vector, commonly known as the kissing bug, releases the parasites into the host. Tissue damage caused by the feeding or the host scratching the bite site allows the parasites to enter the bloodstream and disseminate throughout the body, primarily targeting nucleated cells. Once inside, the parasite multiplies and undergoes differentiation processes until the host cell ruptures, releasing new parasites into the bloodstream to continue the cycle. (b) Infection interface: A key protein involved in this interaction is gp82, which binds to LAMP2. Both gp82 and LAMP2 are glycosylated; gp82 is anchored to the membrane via a GPI anchor, while LAMP2 is embedded in the membrane by a transmembrane domain. This interaction triggers a calcium cascade within the lysosome through PKC activation, leading to F-actin disruption and membrane rupture, facilitating parasite entry into the cell.

A key factor in *T. cruzi*’s evasion strategy is the glycoprotein gp82,^13^ which is expressed on MT surfaces and anchored by glycosylphosphatidylinositol (GPI). Gp82 binds to the host receptor lysosome-associated membrane protein 2 (LAMP2), triggering a cascade of events, including phospholipase C (PLC) activation and the production of diacylglycerol (DAG) and inositol 1,4,5-triphosphate (IP3).^14^ This signaling activates protein kinase C (PKC), leading to an increase in intracellular Ca^2+^, which disrupts the actin cytoskeleton and facilitates parasite internalization (**Fig.1b**).^15^ Additionally, this process recruits lysosomes to the cell periphery, enhancing the parasite’s ability to invade and establish infection.^16^

The interaction between *T. cruzi* and host cells is a complex process, and the recent identification of the gp82 receptor offers promising insights into this interaction.^17^ Despite extensive research on gp82 and LAMP2, several aspects remain unclear, particularly regarding the roles of glycans and membranes in modulating host-pathogen dynamics. ^16,18,19^ Glycosylation is crucial for maintaining the structural integrity and stability of protein interactions and plays a significant role in immune recognition and evasion strategies. ^20–22^ Understanding the role of glycans in the gp82:LAMP2 complex during the infection process is therefore essential. Moreover, elucidating the atomic details of *T. cruzi* invasion mechanisms could lead to new therapeutic strategies, especially given the limitations of current treatments for Chagas disease, such as nifurtimox and benznidazole, which have high toxicity and low patient adherence.^23^ Similar to other host-pathogen interactions, such as the SARS-CoV-2–ACE2 interface where mutations enhance binding a”nity and force stability under mechanical stress, ^24,25^ *T. cruzi* may utilize the gp82 glycoprotein to optimize its attachment and invasion through interactions with LAMP2. This concept aligns with studies of other pathogens, such as reovirus,^26^ where initial weak interactions are strengthened by conformational changes to facilitate infection, and the extreme mechanostability observed in staphylococcal adhesins binding to host fibrinogen.^27–29^ Insights from these studies suggest that a detailed understanding of the gp82:LAMP2 interaction, including its glycan components and mechanostability, could uncover novel targets for therapeutic intervention against Chagas disease.

In this study, we provide the first atomistic exploration of the full gp82:LAMP2 complex to elucidate the molecular mechanisms underlying *T. cruzi* infection. We carefully constructed the entire complex at atomic resolution, starting with the 3D structures of gp82 and LAMP2 provided by Onofre et al.^18^ To model the transmembrane region (TM) of LAMP2 (residues 378–410), we employed secondary structure prediction tools, particularly the protein structure prediction software AlphaFold 2.2.0.^30^ This model was further refined through molecular dynamics (MD) simulations to perform a geometry optimization as well as a thermal equilibration of the TM region, utilizing the high-performance capabilities of NAMD 3.^31,32^ To accurately represent the glycosylation states and GPI anchoring of gp82 and LAMP2, we employed the CHARMM-GUI server,^33^ which allowed us to introduce the GPI anchor to gp82 and glycosylate both proteins, as well as to construct biologically relevant membrane environments specific to *T. cruzi* and *H. sapiens*. Using VMD^34,35^ and its QwikMD plugin,^36^ we assembled a comprehensive model comprising the glycosylated gp82:LAMP2 complex, the human membrane, and the *T. cruzi* membrane in an integrated system using VMD 1.9.4. Leveraging the latest advances in computing,^37^ we integrate GPU-accelerated simulations with experimental data to unravel biomolecular interactions at an unprecedented resolution, offering new insights into complex molecular mechanisms and guiding future therapeutic developments. Our approach integrates cutting-edge computational techniques with existing experimental data to provide a comprehensive molecular view of the gp82:LAMP2 interaction. By bridging experimental findings with high-resolution simulations, we provide new insights into the mechanistic underpinnings of this critical step in *T. cruzi* infection, potentially guiding the development of targeted therapeutic interventions.

## Results

### Complex Assembly

To investigate the gp82:LAMP2 interaction in *T. cruzi* infection, we refined the 3D model of the complex provided by Onofre et al. (2021) by adding the missing TM region of LAMP2_378-410_. First, CHARMM-GUI’s Membrane Builder^38^ was employed to construct organism-specific membranes for *T. cruzi* and *H. sapiens* based on known lipid compositions (see **Fig. 2a** and **Table S1**). Since lipid membranes play a crucial role in providing a hydrophilic/hydrophobic barrier,^39^ significantly affecting polarizability^40,41^ and molecular permeation,^42,43^ accurately modeling them to closely resemble real membranes is extremely important. The *T. cruzi* membrane was modeled to reflect its adaptive lipid composition, particularly its high content of phosphatidylcholine (PC),^44–46^ while the human membrane composition was informed by models involving glycans in viral infection processes.^47^ Both membranes were equilibrated using NAMD 3 with restraints on phosphate atoms and dihedrals to ensure proper system stability.

**Figure 2:**
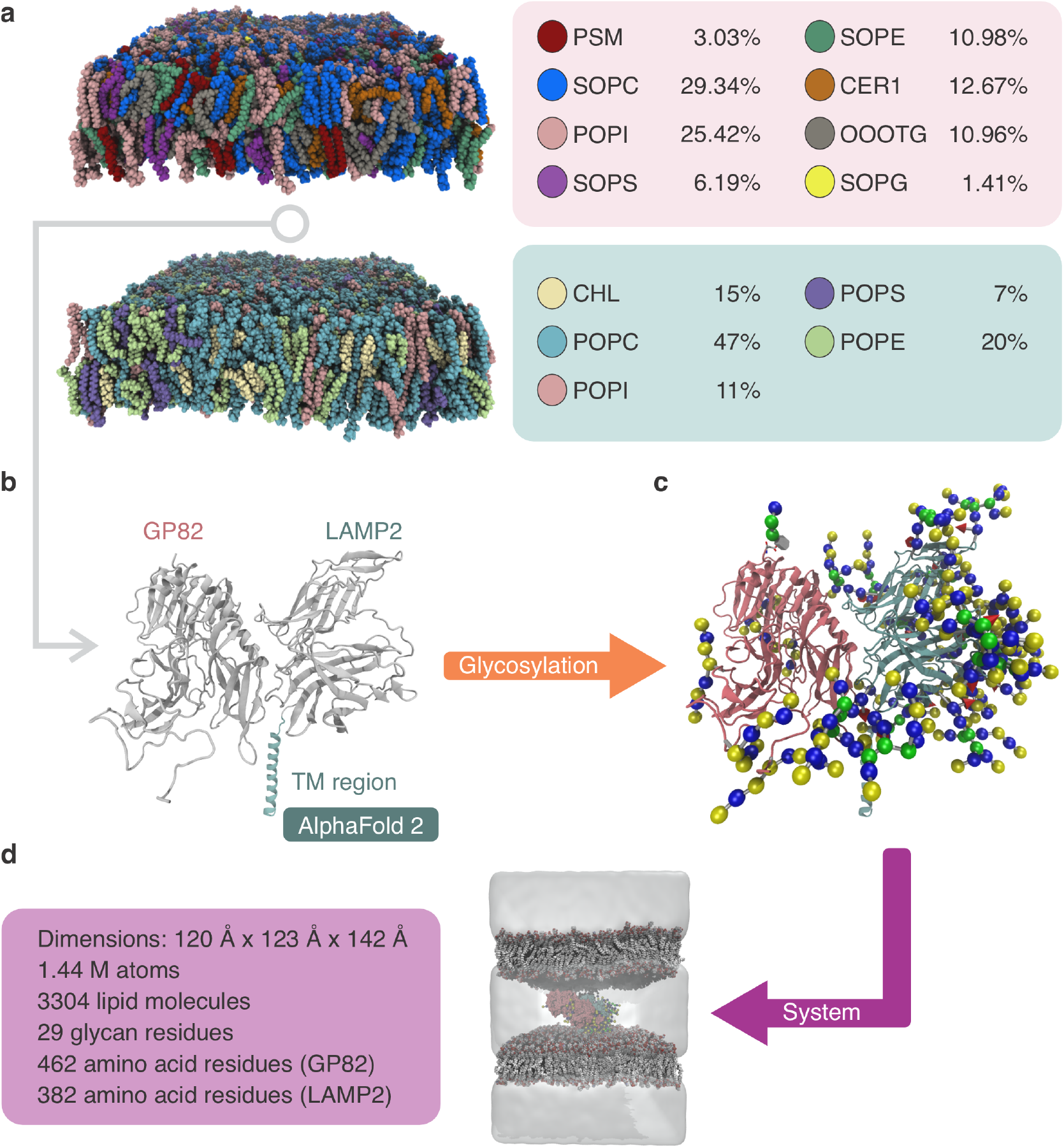
Complex assembly. The rational design of the gp82:LAMP2 complex began with a model from.^18^ (a) The lipidomic constituents of each organism’s membrane were analyzed and modeled using the CHARMM-GUI server for both *T. cruzi* and *H. sapiens*. The transmembrane regions were constructed using AlphaFold to predict the structure accurately. (c) GPI anchors and glycans were designed using CHARMM-GUI to ensure appropriate glycosylation. (d) All components were integrated using VMD, resulting in a complete model comprising 1.44 million atoms. MD simulations were performed using NAMD to stabilize the membrane and protein components.

The TM region of LAMP2, crucial for anchoring this protein to the membrane, was modeled using the Quick2D server for secondary structure prediction^48,49^ and AlphaFold 2.2.0.^30^ Overall, the AlphaFold prediction closely aligned with secondary structure predictions (**Table S2**). We assessed the predicted models using Predicted Alignment Error (PAE) and Local Distance Difference Test (pLDDT) scores, selecting the most reliable structure (Model 3 ipTM+pTM = 0.74) for further study. To integrate the modeled TM region with the existing LAMP2 complex, we used Modeller 10.1 ^50^ to connect it specifically LAMP2_V377_, followed by a 10 ns MD simulation in NAMD 3^31^ to equilibrate the system.

To accurately model the gp82 protein anchored to the membrane via a GPI anchor, we modified the gp82 structure by removing the C-terminal 24 amino acids (GSMHGGVS-RALLLLLGLCGFAALY), allowing for post-translational attachment of the GPI anchor at gp82_D483_ ^51^ (see **Fig. S1**). Both gp82 and LAMP2 undergo extensive glycosylation (**Fig. 2c**), with specific sites for N-linked glycans (**Fig. S2**) on gp82 (S3, T17, T77, S182, T336 ^52^ and multiple N- and O-linked glycosylation (**Fig. S2, S3**) sites on LAMP2 ^53,54^ (see **Table S3**). Using the CHARMM-GUI Glycan Reader and Modeler, we simulated these glycosylations and added the GPI anchor.

With all components assembled and equilibrated using MD simulations, the complete system – comprising the glycosylated gp82(GPI anchor):LAMP2 complex and the corresponding membranes – was assembled using VMD^35^ (**Fig. 2d**). We then performed 50 ns of MD simulations to equilibrate the assembly, followed by 1 *µ*s of production MD simulations, enabling the first comprehensive evaluation of the gp82:LAMP2 complex.

### GP82:LAMP2 Interaction

In light of recent findings, the molecular mechanisms governing the interaction between *T. cruzi* gp82 and host cell LAMP2 during metacyclic trypomastigote invasion raises several unresolved questions. Manque et al. (2000)^55^ identified key binding regions within gp82 that are critical for cell adhesion, yet the minimal set of pairs necessary to stabilize this interaction and drive parasite internalization remains unclear. Onofre et al. (2021) ^18^ identified specific pairs – gp82_E230_ and gp82_K233_ on the parasite, and LAMP2_N120_ and LAMP2_N121_ on humans – as central to this interaction. However, whether these two pairs of pairs alone are su”cient for stable binding, considering the extensive glycosylation of both gp82 and LAMP2, remains an open question.

Furthermore, while GPI anchors and membrane glycans are common features of both gp82 and LAMP2, their role in mediating or stabilizing this interaction has not been investigated. These unexplored factors, along with the precise contribution of membrane components and post-translational modifications, may provide a more comprehensive understanding of how gp82:LAMP2 interactions initiate the signaling cascades that lead to host cell invasion.

To address these questions, using VMD,^34,35^ we analyzed our MD trajectories to evaluate pairs contacts within 4 Å(**Fig. 3a**). The contact analysis revealed key interactions consistent with previous experimental studies, including the well-established pairs gp82_K233_:LAMP2_N120_ and gp82_K233_:LAMP2_D121_, which have been implicated in gp82:LAMP2 binding.^18,55^ Over the course of the simulation, we noticed that gp82_K233_:LAMP2_N120_ was replaced by another contact, namely gp82_K233_:LAMP2_L119_, keeping the same region of interaction but with a different pair. In addition to these known contacts, our simulations identified novel pairs and contact regions, such as gp82_193-194_:LAMP2_331-332_, gp82_R215_:LAMP2_Q331_, gp82_224-227_:LAMP2_330-333_, gp82_T229_:LAMP2_S299_, gp82_S236_:LAMP2_E158_, gp82_K244_:LAMP2_154,156-157_, along with connections in a new region, specifically gp82_R277_:LAMP2_N151_, bringing light to previously uncharacterized interactions between gp82 and LAMP2. While residues gp82_224-227_, gp82_229-230_, gp82_K233_, gp82_K236_, gp82_K244_, gp82_R277_, and LAMP2_120-121_ were noted in experimental work with gp82,^18,55^ this is the first time these interactions with LAMP2 have been described, highlighting potential new insights into the molecular basis of *T. cruzi* invasion.

**Figure 3:**
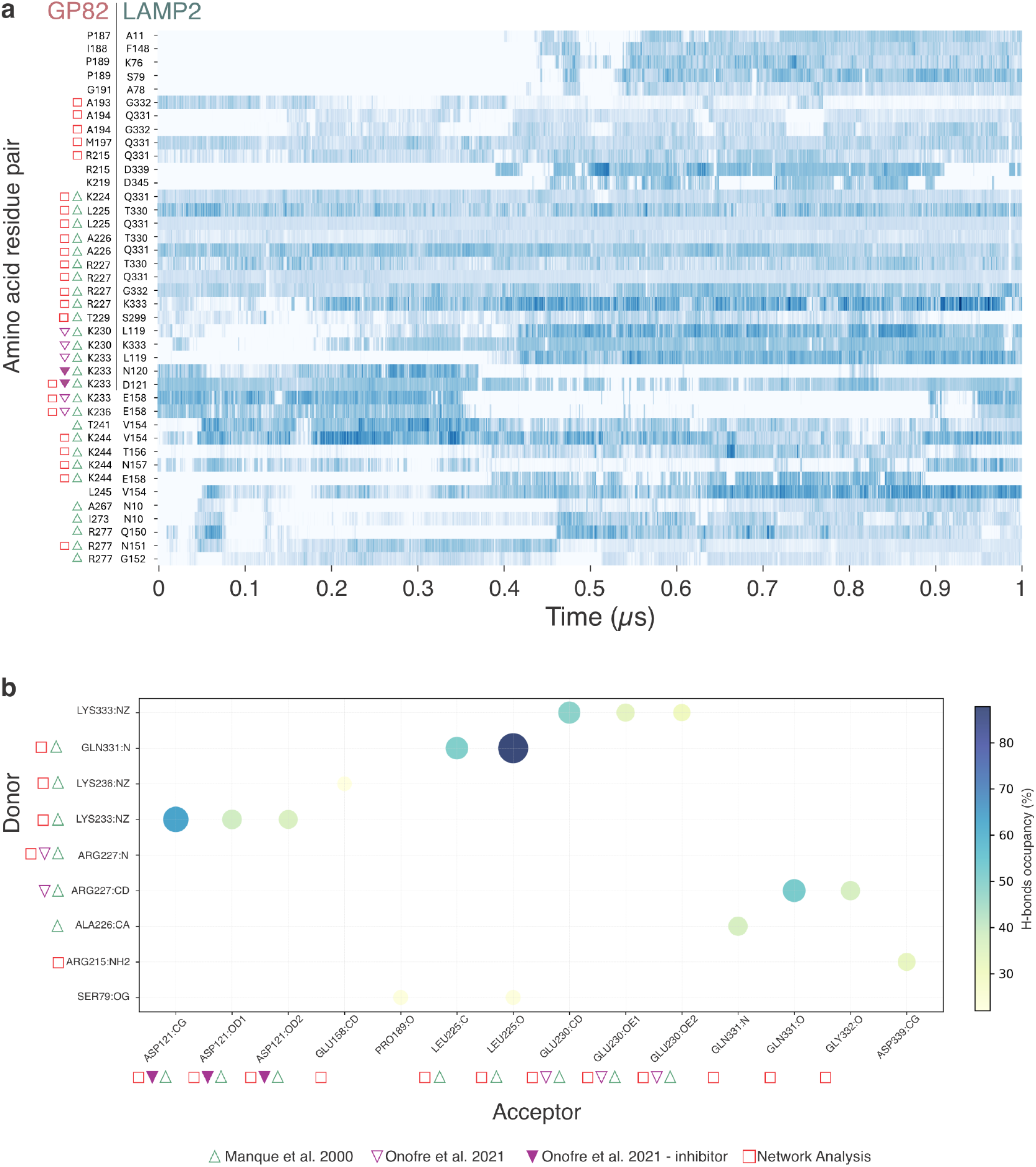
Analysis of contacts and hydrogen bond occupancy during the simulation. (a) Timeline of the number of contacts per residue pair: This plot tracks the temporal evolution of residue-residue contacts throughout the simulation. (b) Scatter plot of hydrogen bond occupancy (*>*20%): Acceptor and donor amino acids are represented on the x- and y-axes, respectively. The color and size of each circle correspond to the hydrogen bond occupancy value. Triangles and squares mark residue pairs identified from experimental data or network analysis, highlighting key interactions.

Additionally, we evaluated hydrogen bonds between gp82 and LAMP2 residues during the MD simulation using VMD. Hydrogen bond occupancy analysis, considering bonds with *>* 20% occupancy, revealed that the most stable hydrogen bond interactions involved previously described residues (**Fig. 3b**). Notably, the highest occupancies were observed for pairs identified in the contact analysis, such as gp82_225-227_:LAMP2_Q331_ and gp82_K233_:LAMP2_D121_, further supporting the critical role of these interactions in the gp82:LAMP2 binding interface. These findings reinforce the importance of these pairs in stabilizing the interaction and suggest a possible mechanism for enhancing parasite invasion.

However, hydrogen bonds are not always the dominant form of interaction in biomolecular interfaces. Recent studies have highlighted the importance of other mechanisms. For example, in bacterial cellulosome assembly, increased contact area helps stabilize protein complexes under mechanical stress.^56,57^ Similarly, non-hydrogen bond contacts and collective motions are crucial for stabilizing interactions in filamins^58,59^ and for the binding of SARS-CoV-2 to its human receptor.^25^ A more powerful approach to identifying critical pairs during binding involves Dynamic Network Analysis, which our group has successfully employed to investigate various protein interactions, including SARS-CoV-2 proteins,^24^ Staphylococci adhesins,^28^ and most recently, a”body interactions with PD-L1. ^60^ In this study, we applied the dynamical network analysis python package^61^ to obtain correlations of motion from our simulation trajectories. In this methodology, the system is represented as a network of nodes, where each node corresponds to an amino acid residue or glycan, and edges are formed between pairs of nodes that remain in contact throughout the simulation (**Fig. 4a**). Contact between nonconsecutive monomers is defined when any heavy atoms (excluding hydrogen) from the two residues are within 4.5Åof each other for at least 75% of the analyzed frames. This network-based approach provides a comprehensive view of the correlated motions within the gp82:LAMP2 complex, helping to pinpoint critical pairs that stabilize the interaction beyond those identified through traditional contact and hydrogen bond analyses.

**Figure 4:**
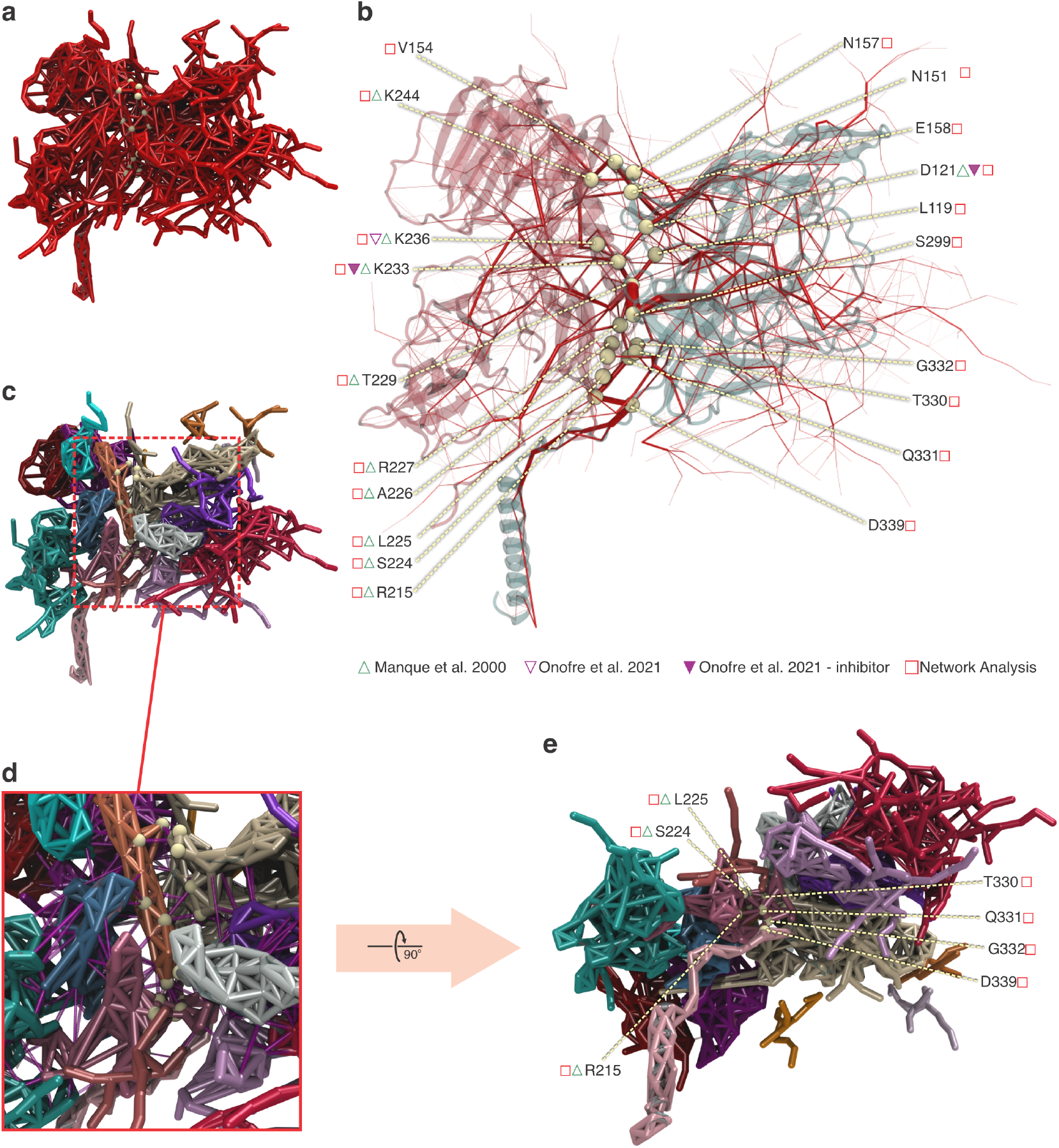
Network analysis of gp82:LAMP2 interactions. (a) gp82:LAMP2 interaction rendered as a generalized correlation-based dynamical network, where edge weights represent generalized correlation coe”cients. (b) Betweenness centrality measurements of the network, highlighting the homogeneous distribution of interactions between key pairs, shown in yellow. Communities represented by different colors in the protein’s secondary structure, with a zoomed-in view showing the pairs involved in the interaction interface (in yellow) (d). (e) Rotation along the X-axis by 90^×^of (c), showing that the gp82_R215_:LAMP2_330–332, 339_ pairs are located within the same community, representing a critical connection to the transmembrane region.

The Dynamic Network Analysis provided valuable insights into the critical pairs mediating the gp82:LAMP2 interaction. Betweenness analysis revealed an edge with a significantly higher betweenness compared to others, which was gp82_T229_:LAMP2_S299_, indicating the importance of these region and its influence across the network (**Fig. 4b**). Betweenness is a measure in network analysis that quantifies the extent to which a node or edge acts as a bridge along the shortest paths between other nodes. High betweenness indicates a critical role in facilitating communication or flow within the network, highlighting its importance in connecting disparate parts of the system. ^61^ Other important pairs were identified, such as gp82_K233_:LAMP2_D121_, gp82_R215_:LAMP2_330-332, 339_, and gp82_K244_:LAMP2_154,156-157_, reflecting the distribution of interactions across the interface and the crucial involvement of gp82_R215_:LAMP2_D339_, connecting the TM region. These interactions were also highlighted when analysing allosteric communication pathways^62^ between the human and parasite membrane. Community analysis further highlighted the cooperative motion between the pairs in the interface (yellow in **Fig. 4c and d**) and the linking part to the TM region (yellow in **Fig. 4e**). Moreover, novel interactions were observed in proximity to experimentally validated residues, including gp82_224-227_:LAMP2_330-332_, gp82_T229_:LAMP2_S299_, gp82_K233_:LAMP2_L119_, gp82_S236_:LAMP2_E158_, gp82_K244_:LAMP2_154,156,157_, in addition to the gp82_K277_:LAMP2_N151_ interaction, suggesting new regions of cooperative motion that could play a previously uncharacterized role in stabilizing the complex (**Fig. 5a**).

**Figure 5:**
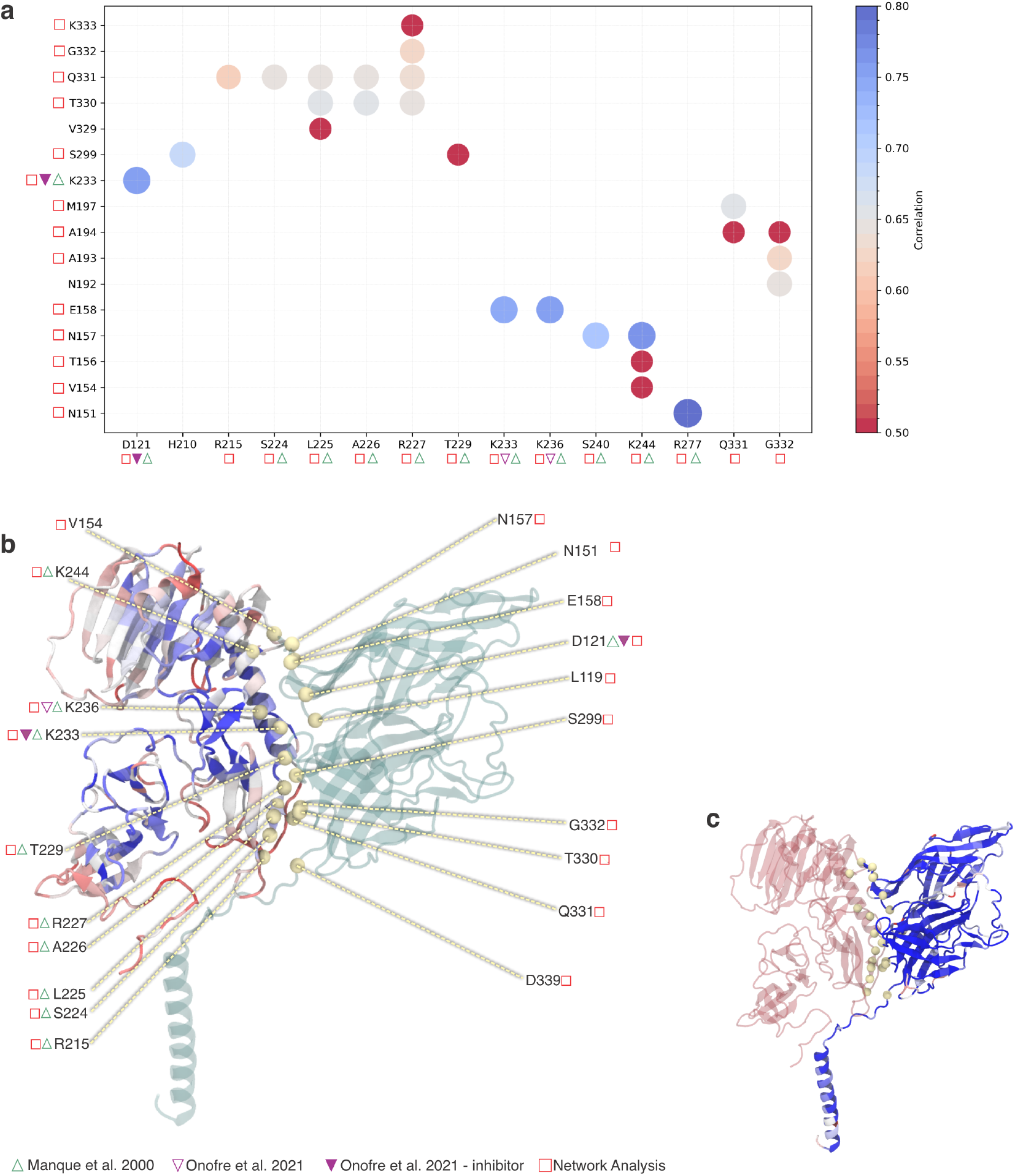
Correlation and conservation analysis of the gp82:LAMP2 complex. (a) Correlation values between residue pairs, with red indicating higher correlation and blue indicating lower correlation. (b) Sequence conservation analysis performed using BlastP, with regions colored by conservation level—blue for highly conserved regions and red for less conserved regions in gp82 and LAMP2 (c). The key interacting pairs are highlighted in yellow, illustrating their functional importance.

The analysis of the molecular dynamics trajectories provided valuable insights into previously uncharacterized interactions between gp82 and LAMP2, revealing new potential contact regions and patterns of correlated motion. These findings suggest novel mechanisms of interaction that extend beyond the critical pairs identified by. ^18,55^ To further evaluate the importance of these new interactions, we compared our results with additional sources of experimental data, specifically genomic sequence databases, to explore how evolutionary conservation supports the significance of these newly identified interactions.

To assess the evolutionary importance of the identified key pairs, we performed a sequence conservation analysis using BlastP. ^63^ This analysis revealed that the key interacting pairs are located in highly conserved regions of both gp82 and LAMP2 (**Fig. 5b,c**). We used the BlastP tool to align sequences for gp82 (GenBank accession number L14824) and LAMP2 (UniProtKB P13473), selecting sequences with at least 30% identity and query coverage. After trimming the sequences, we visualized the regions of conservation using VMD’s Multiseq, where conserved regions were color-coded according to their level of conservation. This visualization confirmed that the key pairs identified in our network and experimental analyses, including gp82_K233_:LAMP2_D121_, gp82_224-227_:LAMP2_330-332_, gp82_T229_:LAMP2_S299_, gp82_K233_:LAMP2_L119_, gp82_K236_:LAMP2_E158_, and gp82_K244_:LAMP2_154,156,157_, are situated in conserved areas, suggesting their critical role in the gp82 interaction across different species (**Fig. 5b**). As expected, the human LAMP2 protein is heavily conserved (**Fig. 5c**).

## Discussion

In this study, we presented the first comprehensive model of the gp82:LAMP2 complex, incorporating key molecular features such as glycans, biologically relevant membrane compositions, and GPI anchors. Our MD simulations and subsequent dynamic network analysis provided detailed insights into the interactions between *T. cruzi* gp82 and the human host receptor LAMP2. We confirmed the importance of previously identified residues, such as gp82_224-227_, gp82_T229_, gp82_K233_, gp82_K244_, gp82_K277_, and LAMP2_D121_, in stabilizing the complex. In addition, we identified novel critical regions of interaction, such as gp82_R215_:LAMP2_331-339_, gp82_K244_:LAMP2_154,156-157_, gp82_K233_:LAMP2_L119_ which had not been described in previous studies. These findings highlight the complexity of the gp82 interaction, suggesting that previously unrecognized pairs may play a crucial role in facilitating the parasite’s invasion of host cells.

Furthermore, to gain deeper insights into the dynamic evolution of the gp82:LAMP2 interaction over the course of the simulation, we performed a network analysis to assess residue correlations and community formation at three distinct time points, representing the early, mid, and late stages of the simulation. Notably, at the beginning of the simulation, the correlation analysis revealed strong interactions between residues such as gp82_K233_:LAMP2_D121_, in agreement with previous experimental studies. These residues displayed high betweenness values, indicating their key roles in maintaining the initial stability of the complex. As the simulation progressed, these strong correlations became more evenly distributed across other regions of the complex, reflecting a more balanced interaction network. Similarly, our community analysis demonstrated that, initially, the TM region of LAMP2 and its main chain formed distinct and separate communities. However, by the end of the simulation, a clearer integration of the TM region with the rest of the LAMP2 chain was observed, suggesting increased connectivity and communication across of the interaction complex S4.

Moreover, while we confirmed the crucial role of the acidic region in gp82 for LAMP2 binding, our analysis also identified a new polar interaction region that may play an equally important role in stabilizing the complex. Interestingly, despite incorporating glycans and membrane lipids into our model, these biomolecules did not appear to significantly influence the interaction between gp82 and LAMP2, at least after the contact is formed. However, the TM region of LAMP2 showed to be important in the stabilization of the interaction interface. Future research could focus on determining whether other components, such as extracellular matrix elements, might have an impact on the interaction dynamics.

Overall, our findings not only confirm key experimental observations but also extend the understanding of the molecular underpinnings of *T. cruzi* invasion, highlighting potential new targets for therapeutic intervention. By identifying novel critical pairs and regions within the gp82:LAMP2 interface, this study provides a foundation for the development of drugs aimed at disrupting this interaction, potentially aiding in the fight against Chagas disease.

## Methods

### Modeling the gp82:LAMP2 complex

The structures of *T. cruzi* gp82 and *H. sapiens* LAMP2 were obtained from Onofre et al. (2021).^18^ However, the provided gp82:LAMP2 complex lacked the TM region of LAMP2 (residues 378-410), which is crucial for studying the full interaction of these proteins. To model this missing region, we first retrieved the LAMP2 sequence from UniProtKB (P13473) and used the Quick2D server^48,49^ to predict secondary structure elements and detect TM regions. This initial step provided a general overview of the domain architecture.

We then utilized AlphaFold 2.2.0^30^ with default settings to predict the 3D structure of the TM region. AlphaFold is known for its high accuracy in predicting protein structures, and its Predicted Aligned Error (PAE) plots and Local Distance Difference Test (pLDDT) scores were used to assess the quality of the models. The model with the lowest PAE and highest pLDDT scores, indicating high structural confidence, was selected. Modeller 10.1^50^ was then used to incorporate the predicted TM region into the existing gp82:LAMP2 complex, ensuring that the overall structural integrity of the complex was maintained.

To model the glycans attached to both gp82 and LAMP2, we employed Glycan Reader & Modeler^33^ through the CHARMM-GUI server.^33^ Glycosylation is essential for proper protein function and stability, and the Glycan Reader & Modeler provides a reliable method to build these carbohydrate structures based on known glycosylation sites and glycan sequences. We input the relevant glycan sequences, glycosylation sites, and structures for both proteins into the server. The module used a template-based approach to predict and model the glycan structures, which were then attached to the appropriate sites on gp82 and LAMP2. Additionally, the GPI anchor for gp82 was modeled using the same approach, ensuring that all necessary post-translational modifications were accurately represented in the final complex structure.

### Membrane Modeling

The Membrane Builder – Bilayer Builder module within the CHARMM-GUI server^38**? ? ? ?**^ was employed to construct the two membranes used in this study, representing the membranes of *T. cruzi* and *H. sapiens*. The lipid compositions for both membranes were based on lipidomics data, ensuring biological relevance to the specific organismal membrane environments. For simplicity and consistency, we maintained identical lipid compositions in both the inner and outer leaflets of the membranes.

The membrane dimensions were set with lateral dimensions of 225 Å in the X and Y directions, large enough to accommodate the gp82:LAMP2 complex and ensure proper interaction with the membrane environment. Along the Z axis, a default water layer thickness of 22.5 Å was used to provide su”cient hydration of the system. The lipid bilayers were solvated by a water box using the TIP3P water model, and a 0.15 M NaCl ion concentration was introduced to simulate physiological conditions, similar to the setup used in the gp82:LAMP2 complex.

Following the definition of these parameters in CHARMM-GUI, the necessary input and configuration files for molecular dynamics simulations were automatically generated. These files ensured proper integration of the membranes with the protein complex, allowing for an accurate and stable simulation environment for both the parasite and host cell membranes.

### Equilibration Molecular Dynamics Simulations

Following previously stablished protocols for membrane simulations,^40,64^ MD simulations were conducted using the CHARMM36 force field^65^ in NAMD 3.^31^ Systems were prepared with the TIP3P water model, and each system was neutralized using a 0.15 M NaCl ion concentration. Simulations were performed in the NPT ensemble (constant pressure and temperature), with temperature maintained at 310 K via Langevin dynamics for both temperature and pressure coupling. The pressure was held constant at 1 bar. Short-range non-bonded interactions were calculated using a distance cut-off of 12 Å, while long-range electrostatic interactions were computed using the particle-mesh Ewald (PME) method.^66^

### Protein Complex Equilibration

The protein complex, including the TM region modeled by AlphaFold, was subjected to energy minimization for 5,000 steps prior to MD simulations. The system was then equilibrated by performing a 2 ns MD simulation in the NPT ensemble, with a timestep of 2.0 fs. During this phase, position restraints were applied to the protein backbone atoms to stabilize the structure. Following this pre-equilibration, the production simulation was run for 20 ns under the same conditions, with a 2.0 fs timestep and restraints removed.

Before conducting Steered Molecular Dynamics (SMD) simulations, the protein complex (gp82:LAMP2) was submitted to an additional energy minimization step of 1,000 steps, this time under the NVT ensemble (constant volume and temperature) with position restraints on the backbone atoms. A temperature ramp was then applied, gradually increasing from 0 K to 310 K over the first 0.5 ns. This was followed by a 1 ns equilibration at 310 K in the NPT ensemble, with a 1.0 fs timestep.

For the SMD simulations, a constant pulling speed of 5 × 10^−4^ Å/ps was applied along the Z-axis, with a spring constant of 5 kcal/mol·Å^2^, over a period of 10 ns, using a 1.0 fs timestep. The C-terminal V329 residue was anchored, and the N-terminal F382 residue in LAMP2 was pulled, aligning the system for coupling with the membrane.

### Membrane Equilibration

Membrane equilibration began with an energy minimization protocol of 10,000 steps. Restraints were then applied to the phosphate atoms in the Z direction and the lipid dihedrals to maintain membrane stability. Successive equilibration steps were performed to equilibrate the simulation, following standard membrane simulation protocols. The system was then simulated under the general simulation conditions, with a timestep of 2.0 fs for a total of 10 ns. The protocol was repeated for both membranes.

### System Assembly

After equilibrating all individual components through MD simulations, the complete system—consisting of the glycosylated gp82 protein with its GPI anchor, LAMP2, and the corresponding lipid bilayer membranes—was assembled using the Visual Molecular Dynamics (VMD) software.^35^ The gp82_GPI_ anchor and LAMP2 were carefully positioned in their respective membrane environments, ensuring proper embedding of LAMP2’s TM region within the lipid bilayer and the insertion of the GPI-anchored gp82 into the membrane of *T. cruzi*. The complex was solvated using the TIP3 water model,^67^ and the net charge of the protein was neutralized using a 0.15 M salt concentration of sodium chloride. The total system measured 120 Å × 123 Å × 142 Å and contained 1.44 million atoms. The final assembly included 3,304 lipid molecules, two proteins — gp82 with 462 amino acid residues and LAMP2 with 382 residues — both glycosylated with 29 glycan. This setup allowed for the first detailed, fully atomistic investigation of the gp82 complex in a biologically relevant membrane environment.

### Production Molecular Dynamics Simulations

The fully assembled system was subjected to all-atom molecular dynamics simulations using NAMD 3^31^ with the CHARMM36^65^ force field, following QwikMD standard protocols.^36^ To maintain structural integrity during equilibration, harmonic restraints were applied to the proteins. All MD simulations were conducted under periodic boundary conditions, with a Å cut-off applied to short-range non-bonded interactions, while long-range electrostatics were treated using the PME method.^66^ Prior to the MD simulations, the system underwent 1,000 steps of energy minimization. Subsequently, a 50 ns MD simulation with positional restraints on protein backbone atoms was performed, with temperature ramping from 0 K to 310 K over the first 1.0 ns, using a 2.0 fs timestep in the NVT ensemble to pre-equilibrate the system. Production MD simulations were conducted in the NPT ensemble at a constant temperature of 310 K, regulated via Langevin dynamics, and pressure maintained at 1 bar. A timestep of 4 fs was used for all production runs, enabled by hydrogen mass repartitioning. ^68^ The total duration of the production MD simulations was 1 *µ*s. Simulations were performed using Cybershuttle cyberinfrastructure.^69^

### Analysis

The simulations were analyzed using in-house Python and TCL scripts along with VMD.^34,35^ Contact maps for residue-residue interactions were generated with VMD and PyContact,^70^ while hydrogen bonds were also calculated using VMD. Network analysis, performed with the Dynamic Network Analysis package,^61,71^ identified key communication pathways within the protein complex. Force propagation profiles were mapped using Onsager’s reciprocal relations following Schoeler et al. 2015. ^72^ Residue conservation was evaluated via BLOSUM matrices in VMD’s MultiSeq tool, highlighting functionally important pairs. Together, these analyses provided a comprehensive view of the system’s structural and dynamic properties.

## Acknowledgement

RCB acknowledges support from the National Science Foundation (Grant MCB-2143787) and the National Institute of General Medical Sciences (NIGMS) of the NIH (Grant R24-GM145965). RSLR and MLS were funded by the Brazilian Federal Agency for Support and Evaluation of Graduate Education (CAPES) through the CAPES-FIOCRUZ/PrInt program (N 01/2022) and by National Council for Scientific and Technological Development/CNPq/Brazil. Computational resources were provided by Auburn University through start-up funds for RCB.

