## Supplementary Material for "Atomistic Insights into gp82 Binding: A Microsecond, Million-Atom Exploration of Trypanosoma cruzi Host-Cell Invasion"

**List of Associated Content:**

**Supporting Information Tables:**

- **Table S1 - Membrane Composition.**
- **Table S2 - Modeling validation for TM region of LAMP2.**
- **Table S3 - Glycans Composition.**

**Supporting Information Figures:**

- **Figure S1 - Glycosylation Scheme - GPI anchor.**
- **Figure S2 - Glycosylation Scheme - N-glycans.**
- **Figure S3 - Glycosylation Scheme - O-glycans.**
- **Figure S4 - Network Analysis Over Time.**

Table S1: Membrane Composition

| **Lipid type** | **Abbr.** | ***Trypanosoma cruzi*** | |
|---|---|---|---|
| | | **%** | **N** |
| Ceramide | CER160 | 11.95% | 193 |
| Phosphatidylcholine | SOPC | 29.34% | 517 |
| Phosphatidylethanolamine | SOPE | 10.98% | 174 |
| Phosphatidylglycerol | SOPG | 1.41% | 13 |
| Phosphatidylinositol | POPI | 25.42% | 445 |
| Phosphatidylserine | SOPS | 6.19% | 104 |
| Sphingomyelin | PSM | 3.03% | 37 |
| Triacylglycerol | OOOTG | 10.96% | 176 |
| **Total number** | | 1659 | |
| **Average Area** | | 237.65Å | |
| **Lipid type** | **Abbr.** | ***Homo sapiens*** | |
| | | **%** | **N** |
| Phosphatidylcholine | POPC | 47% | 812 |
| Phosphatidylethanolamine | POPE | 20% | 330 |
| Phosphatidylinositol | POPI | 11% | 168 |
| Phosphatidylserine | POPS | 7% | 95 |
| Cholesterol | | 15% | 240 |
| **Total number** | | 1645 | |
| **Average Area** | | 235.27Å | |

Table S2: Modeling validation for TM region of LAMP2.

| LAMP2 TM region | PAE | pLDDT | ipTM + pTM |
|---|---|---|---|
| **Model 1** | 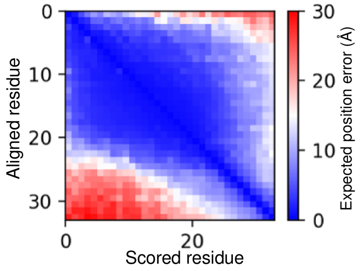 | 79.6 | 0.62 |
| **Model 2** | 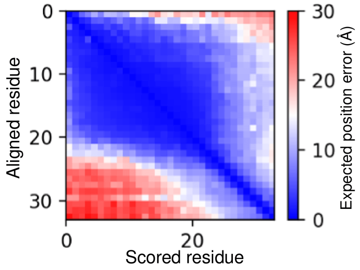 | 78.7 | 0.63 |
| **Model 3** | 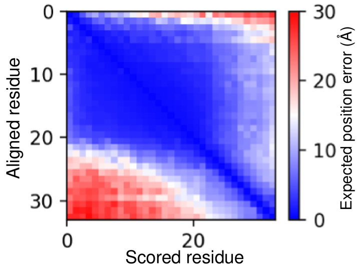 | 81.5 | 0.74 |
| **Model 4** | 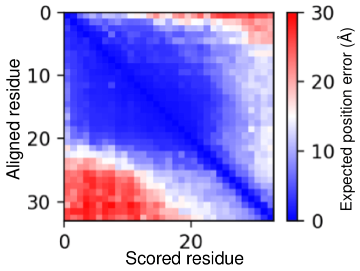 | 81.6 | 0.62 |
| **Model 5** | 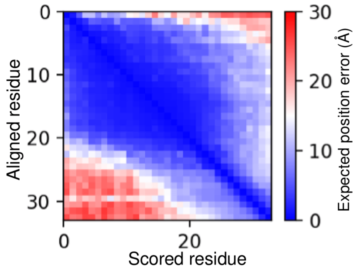 | 80.9 | 0.60 |

Table S3: Glycans Composition.

| Protein | Site | Composition and Type |
|---|---|---|
| **gp82** | D483 | GlcN(1)Hex(3) Glycosylphosphatidylinositol (GPI) anchor |
| | S3, T17, T177, S182, T336 | HexNAc(4)Hex(4) O-glycans |
| **LAMP2** | N21, N30, N47, N73, N95, N201, N214, N247, N272, N279, N289, N328 | HexNAc(7)Hex(8) Fuc(1) N-glycans |
| | S167, T168, T172, T175, T176, S179, T181, T182, T183, T185 | HexNAc(4)Hex(4) O-glycans |

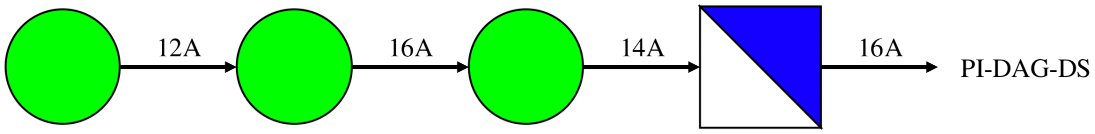


Figure S1: **Glycosylation Scheme - GPI anchor.** Schematic representation of the GlcN(1)Hex(3) Glycosylphosphatidylinositol (GPI) anchor. Blue square - GlcNAc; Green circle - Mannose.

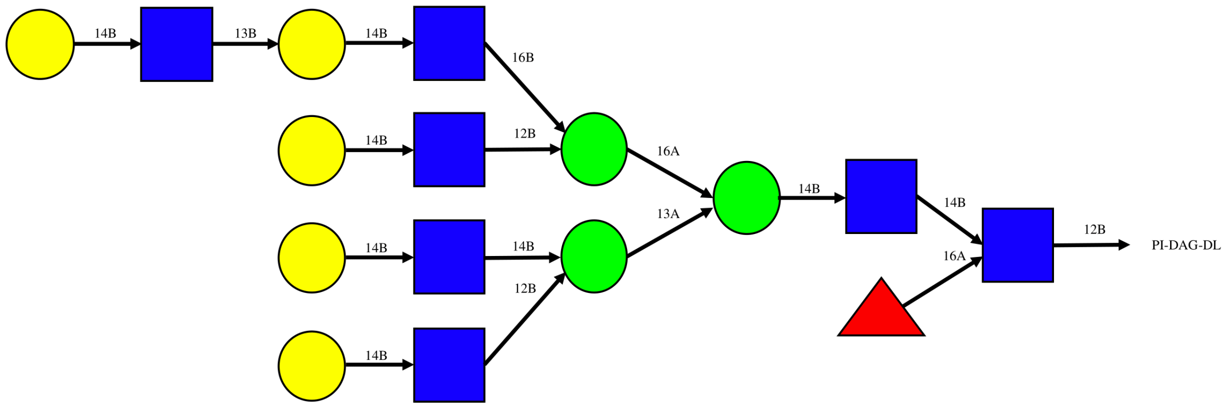


Figure S2: **Glycosylation Scheme - N-glycans.** Schematic representation of the HexNAc(7)Hex(8)Fuc(1) N-glycans. Blue square - GlcNAc; Red triangle - Fucose; Yellow circle - Galactose; Green circle - Mannose.

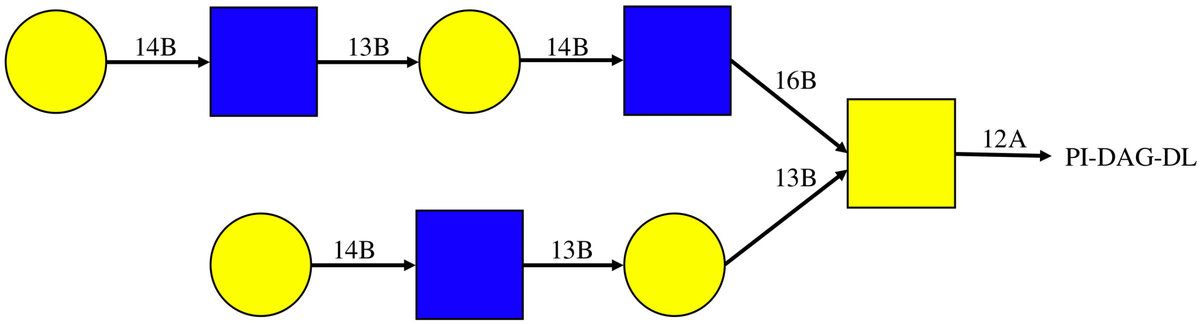


Figure S3: **Glycosylation Scheme - O-glycans.** Schematic representation of the HexNAc(4)Hex(4) O-glycans. Blue square - GlcNAc; Yellow circle - Galactose; Yellow square - GalNAc.

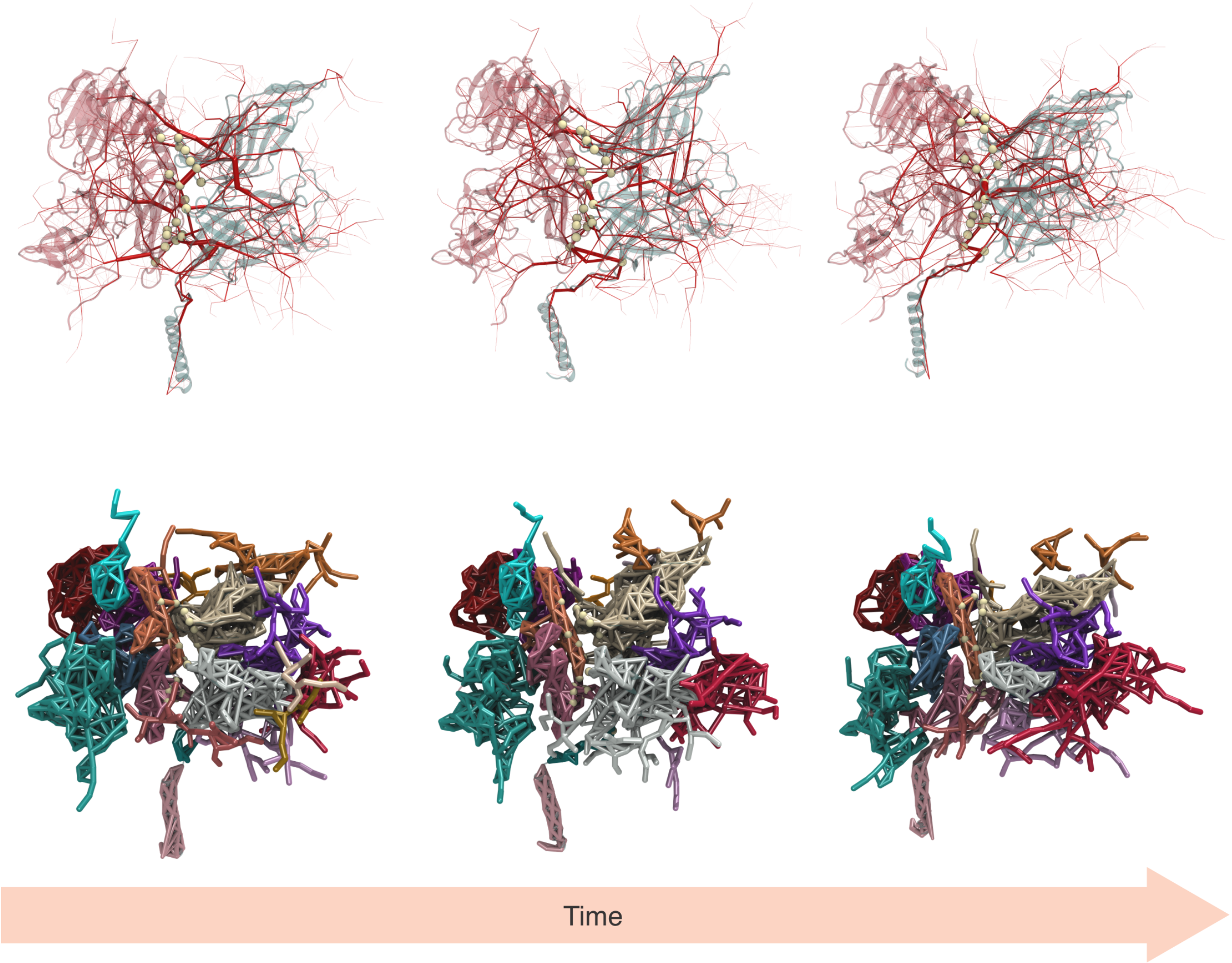


Figure S4: **Network Analysis Over Time.** The first row shows the betweenness analysis, and the second row presents the communities analysis at three distinct stages of the simulation, highlighting key pairs (in yellow) involved in maintaining the stability of the complex.
